# Promising antimicrobial activity of *Moringa oleifera* seed extract fractions

**DOI:** 10.64898/2026.03.23.713792

**Authors:** Godfred Futagbi, Mary Tiwaa Donkor, Bright Churchill Obeng, Stephen Achou, Emmanuella Adjah-Tetteh, Monia Enyonam Honyo, Mary-Magdalene Osei, Selorme Adukpo, Mary Anti Chama, Linda Eva Amoah, Eric Sampane Donkor

**Author notes:** Correspondence author (GF).

## Abstract

This study investigated the antiplasmodial and antibacterial activities of fractionated extracts of *Moringa oleifera* seeds, focusing on the influence of solvent polarity on bioactivity. The results revealed a polarity-dependent distribution of activity. Polar aqueous extracts (crude and residual fractions) exhibited the most pronounced antiplasmodial effects against *Plasmodium falciparum* (3D7 strain), with IC₅₀ values of 107–135 µg/mL. Time-dependent analyses of the crude and residual fractions showed that parasitaemia declined steadily over time, and consequently, percentage inhibition increased with time, with both extracts reaching 70–80% inhibition by 48 hours at higher concentrations. In contrast, moderately polar organic fractions, notably ethyl acetate and dichloromethane, demonstrated strong antibacterial activity against both Gram-positive and Gram-negative clinical isolates, including resistant strains such as MRSA and ESBL-producing Escherichia coli. Minimum inhibitory concentrations (MICs) ranged from 6.3 to 25 mg/mL for the ethyl acetate fraction, and all active fractions exhibited bactericidal properties (MBC/MIC ≤ 4). Comparative analysis showed that while antiplasmodial activity was moderate relative to the standard drug (chloroquine), antibacterial activity was robust and clinically promising. Fractionation revealed that distinct phytochemical classes underlie the two activities: polar compounds appear responsible for antiplasmodial effects, whereas moderately polar compounds drive antibacterial potency. The moderate antiplasmodial activity is significant in the context of adjunctive therapy and resistance management, while the strong antibacterial activity is highly relevant to the global challenge of antimicrobial resistance. Together, the results position *Moringa oleifera* as a promising source of phytochemicals for integrated infectious disease management, particularly in regions where malaria and bacterial co-infections are prevalent.

## Introduction

Despite the implementation of various control and preventive strategies, including the use of insecticide-treated nets (ITNs), indoor residual spraying (IRS), seasonal malaria chemoprevention (SMC), artemisinin-based combination therapies (ACTs) [1], and recently recommended malaria vaccines for children [2], malaria remains a significant cause of illness and death in tropical regions, notwithstanding the success achieved by these interventions. According to global estimates, 282 million malaria cases and 610,000 malaria-related deaths occurred in 2024, indicating an increase over the 2023 figure of 263 million cases and 597,000 deaths. It is a leading cause of morbidity and mortality in sub-Saharan Africa. The WHO African Region bears a disproportionately high burden of malaria, estimated to account for 94% of cases and 95% of deaths [3]. Groups at higher risk of severe infection include children under five, pregnant women and girls, and individuals with HIV or AIDS [3]. Chemotherapy, which is one of the mainstays of malaria control and prevention, faces a significant threat from emerging resistance to antimalarial drugs, including artemisinin [4]. Artemisinin-based combination therapies (ACTs) are currently the most effective treatment for uncomplicated malaria. However, resistance to ACTs has been reported in parts of Southeast Asia and Africa [4,5]. The development of resistance to chloroquine and ACTs poses substantial challenges to effective treatment and control. The emergence of drug-resistant strains of *Plasmodium falciparum* has intensified the search for alternative therapeutics, particularly from plant sources. About 20% of patients in many countries take traditional botanicals to cure malaria [6].

Bacterial infections, which sometimes occur with malaria, are also among the leading causes of ailments and deaths worldwide [7]. The misuse and abuse of antibiotics to manage these infections has led to the development of resistance and the spread of multidrug-resistant (MDR) strains of microbes [8,9]. The rising threat of antimicrobial resistance has necessitated the quest for new antimicrobials.

Plants are known for their medicinal properties and have been used in traditional medicine since antiquity. *Moringa oleifera*, a widely used medicinal and edible plant, is believed to possess bioactive compounds with both antibacterial and antimalarial potential [10–12]. It is also employed as a remedy for some non-infectious diseases, including diabetes, arthritis, and hypertension [11,13], further highlighting its potential.

Furthermore, *M. oleifera* is known not only for its medicinal value but also for its nutritional components. It contains essential minerals, proteins, vitamins, carotene, and amino acids [14; 15]. Its leaves are commonly used as food in many parts of the world [16]. Many parts of the plant are used for treating various diseases, but the leaves are the dominant part used. Therefore, more work has been done on the antiplasmodial activity of the leaves than the other parts, including the seeds. According to Bezerra et al., the leaves accounted for 63% of the parts used, while the seeds accounted for 13%. The leaves are known to be used to treat malaria [11,12,17], and some studies have shown antimalarial potential. It has been noted that *M. oleifera* seeds have been used for treating malaria in many countries [12,18,19], and just like the leaves, the seeds have shown antiplasmodial potential in some studies [20–22]. However, the antiplasmodial properties of *M. oleifera*, particularly the leaves and seeds, and their effectiveness in treating malaria are yet to be fully unravelled. Moreover, studies have shown that the polarity of the solvent used in extraction can affect antimicrobial activity or bioactivity of the extract.

Recent studies have also demonstrated that aqueous and organic solvent extracts of *M. oleifera* seeds and leaves exhibit inhibitory effects on both enteric and non-enteric microbes such as *E. coli*, *Staphylococcus aureus*, *Streptococcus mutans*, *Staphylococcus epidermidis*, *Pseudomonas aeruginosa*, *Proteus vulgaris*, *Enterococcus faecalis*, *Enterococcus cloacae*, *Vibrio cholerae*, *Klebsiella* species, *Trichoderma* species, *Aspergillus flavus*, *Bacillus cereus*, *Streptococcus pneumoniae*, and *Candida* species, among others [11,23,24]. Our previous study has shown that among the extracts examined, aqueous and ethyl acetate leaf and seed extracts were the most effective against both Gram-positive and Gram-negative bacteria, including drug-resistant strains [25].

Although compelling evidence supports both antiplasmodial and antibacterial properties of *Moringa oleifera*, contradictions exist due to differences in extraction methods, plant parts studied, and microbial strains tested [26]. The polarity of solvents significantly influences bioactivity, and fractionation may help isolate beneficial compounds while minimizing undesirable effects. This dual potential highlights *Moringa oleifera* as a promising candidate for further research in combating malaria and bacterial infections simultaneously.

## Materials and methods

### Sampling and preparation of *Moringa oleifera* seeds for extraction

Mature seeds of *Moringa oleifera* were collected from the University of Ghana campus (GPS: GA-522-3371) and surrounding areas, chosen specifically for their lack of chemical pollutants. The seeds were extracted from their pods and dehusked. To preserve the integrity of bioactive compounds, the seeds were shade-dried for two weeks at ambient room temperature. Once dried, the white seed kernels were powdered using a blender.

The powder was stored in clean, sterile, airtight containers until further processing. Botanical identification of the plant material was confirmed by a botanist at the Ghana Herbarium, and a voucher specimen (GF 1603/23) was archived for reference.

### Maceration and extraction of *Moringa oleifera* seeds

Crude extracts of *M. oleifera* seeds were obtained using four solvents, namely ethyl acetate, petroleum ether, absolute ethanol, and distilled water. Following a previously established protocol [25], powdered seeds were soaked in each solvent at a ratio of 1:5 (w/v), consistent with the standard maceration range (1:4 to 1:16) [27]. The seed-solvent mixtures were left to stand for 72 hours to allow thorough extraction of phytochemicals. This process was repeated multiple times by reapplying fresh solvent to the remaining residues. The crude extracts obtained were then filtered using Whatman filter paper to remove suspended insoluble particles. Organic solvent extracts were concentrated using a rotary evaporator (Cole-Parmer), while aqueous extracts were freeze-dried.

### Fractionation of crude aqueous extract

Analysis of the crude extracts showed that the aqueous extract was more potent than the organic extracts. Therefore, the aqueous extract was successively extracted to obtain fractions. Briefly, after maceration of seeds in cold distilled water, followed by filtration using Whatman filter paper, as described above, the resulting aqueous extract was divided into two portions: one was frozen, and the other was freeze-dried. To concentrate the extract for fractionation, the freeze-dried seed extract was dissolved in a portion of the extract that was not freeze-dried and stirred until fully dissolved. This concentrated solution was then subjected to successive maceration using organic solvents: hexane, dichloromethane, ethyl acetate, and butanol, in that order. Each solvent produced a specific fraction, namely hexane, dichloromethane, ethyl acetate, butanol fractions, and a residual aqueous fraction (Fig 1). The fractionation process was repeated four times for each solvent over 24 hours, using a total volume of approximately 3 litres per solvent. All collected organic fractions were concentrated and dried using a rotary evaporator. The residual aqueous fraction was freeze-dried and stored at −20 °C.

**Fig 1.**
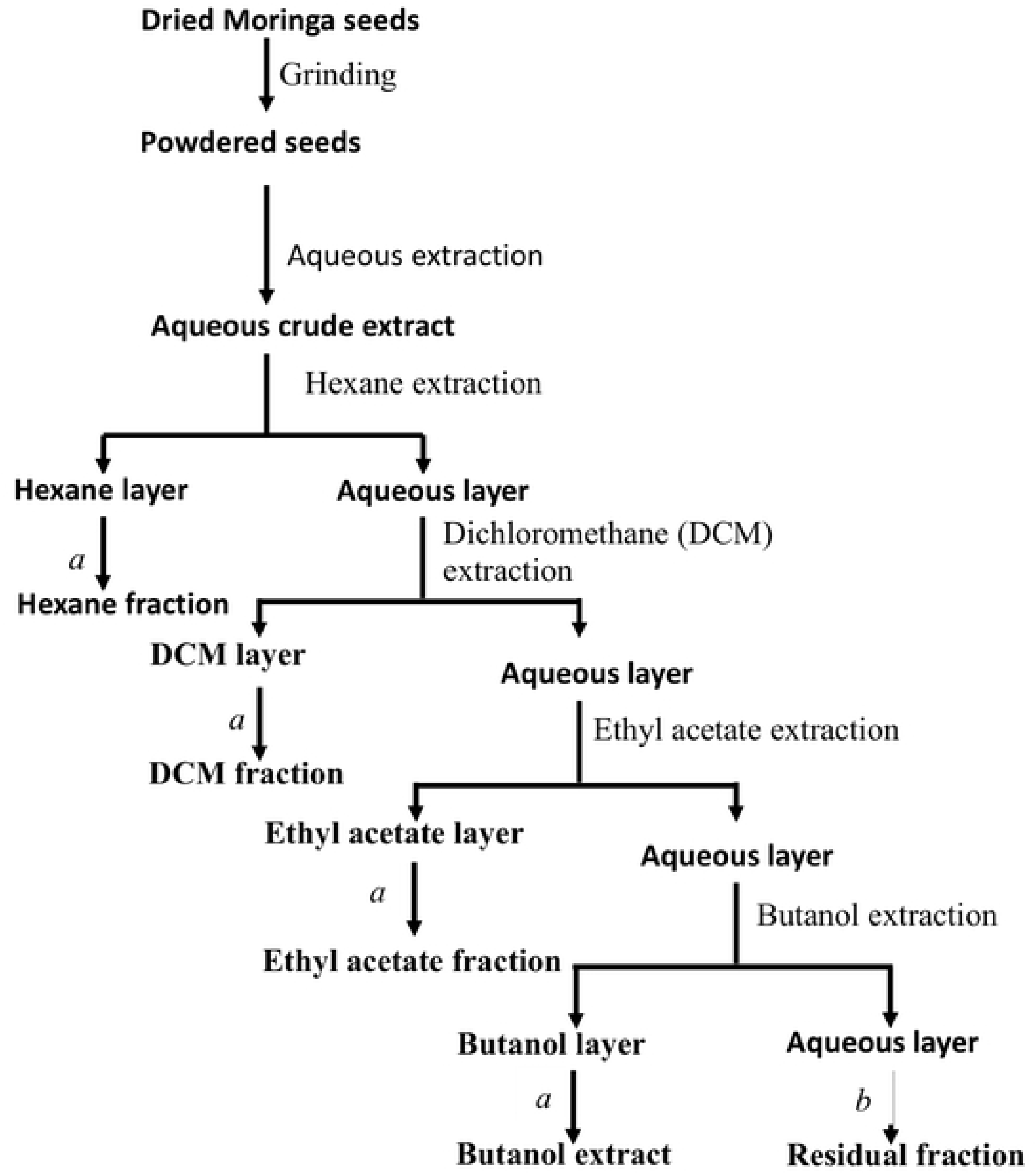
Schematic diagram of the extraction and fractionation procedure. a and b denote rotary evaporation and freeze-drying, respectively.

### Culturing of asexual *Plasmodium falciparum*

Continuous *in vitro* cultures of the chloroquine-sensitive 3D7 strain of *Plasmodium falciparum* were maintained in complete parasite medium (CPM), composed of RPMI 1640 (Gibco, UK) supplemented with 25 mM HEPES, 2 mM L-glutamine, 0.5% Albumax II, 0.5% sodium bicarbonate (NaHCO₃), 50 µg/ml glucose, 50 µg/ml gentamycin and 0.367 mM hypoxanthine. Parasites were cultured in O⁺ human red blood cells (RBCs) and incubated at 37 °C under a gas mixture of 92.5% nitrogen, 5.5% carbon dioxide, and 2% oxygen. Media were changed daily to maintain optimal growth conditions. Once parasitaemia reached ≥5% ring-stage parasites, cultures were synchronized using 5% sorbitol treatment. Cultures were incubated for 48 hours after synchronization to allow ring-stage development. Parasitaemia was then adjusted to 1% and haematocrit to 4% before use in downstream assays.

### Dose-dependent analysis of antiplasmodial activity

The antiplasmodial activity of each extract was assessed by SYBR Green method. Briefly, each of the extracts was dissolved in DMSO except the aqueous extract and residual fraction, which were dissolved in dH_2_O (double distilled water) to obtain an initial concentration of 200 ug/ml. Fifty microliters (50 µL) of each extract were plated in eight concentrations using a two-fold serial dilution (200 µg/mL to 1.56 µg/mL), starting at 200 µg/mL. Each concentration was plated in triplicate. To each treated well, 50 µL of synchronized asexual *P. falciparum* culture at 1% parasitaemia was added. After the plates were placed in a modular incubation chamber, they were gassed for 6 minutes to maintain optimum conditions. After gassing, the chamber was transferred to a 37 °C incubator for 72 hours. The culture was thawed at room temperature after an overnight freezing of the plates followed by the addition of 100 µL of lysis buffer supplemented with SYBR Green, to each well. The plates were incubated for 1 hour and 30 minutes at room temperature, then read using a Varioskan fluorometer at excitation and emission wavelengths of 497 nm and 530 nm, respectively, to obtain the relative fluorescence units (RFU).

### Time-dependent analysis of antiplasmodial activity

Crude extract and residual fraction of *M. oleifera* seed that exhibited moderate activity were tested for their ability to inhibit the growth of *P. falciparum* over time. Each extract was prepared at IC₅₀ and 2 **×** IC₅₀, plated in triplicate, and incubated under the conditions described above. At multiple time points (6, 12, 24, 36, and 48 hours), thin blood smears were prepared and stained to assess parasitaemia levels microscopically.

### Identification, culturing and preparation of bacteria isolates

Pure clinical isolates, control isolates and drug-resistant isolates of gram-negative (*Escherichia coli*, *Pseudomonas aeruginosa*) and gram-positive (*Staphylococcus aureus*) bacteria were obtained from the Department of Medical Microbiology, University of Ghana Medical School. The gram-negative bacteria were cultured on MacConkey agar and the gram-positive bacteria on Mannitol salt agar using standard methods and then incubated at 35 ± 2°C for 24 hours. The cultured bacteria on the MacConkey agars and Mannitol salt agars were picked and separately inoculated into 5ml sterile saline contained in bijou bottles. The turbidity was measured to meet the standard McFarland turbidity at a measurement range of 0.5-0.6 McFarland unit.

### Antibacterial Assay: Agar well diffusion method

The most common methodology for assessing the bioactive chemicals’ antibacterial activity, the agar well diffusion test, was employed. Culture media (Mueller Hinton agar) were prepared according to the manufacturer’s instructions. To confirm the sterility of the agar plates prepared, they were incubated at 37°C for 24 hours and examined for the growth of any microbes. The bacteria were evenly spread on the entire surface of the agar plate. Wells of diameter of five millimetres were made in the agar and a drop of the extract fractions at concentrations of 400 µg/mL and controls were added to the wells. Antibiotics, Amikacin and Linezolid were used as positive controls for gram-negative and gram-positive bacteria, respectively, at 30 µg/ml. After incubation at 37°C for 24 hours, the antibacterial activity was evaluated by the measuring the diameter of the inhibition zone around the wells. This was done for a standard, two drug-resistant and eight clinical isolates each of *Escherichia coli* (from urine and blood)*, Pseudomonas aeruginosa* (from blood) and *Staphylococcus aureus* (from skin lesions).

### Determination minimum inhibitory concentration (MIC)

The minimum inhibitory concentrations (MIC) of the fractions with the best activity, ethyl acetate and dichloromethane fractions, and crude extract were determined using the broth dilution method. Two-fold dilutions of extract fractions were made from the 400 µg/ml concentration of the fractions. With the 96 well plates and according to the inoculum standardization, 100µl of the bacterial isolates were added to 100µl of the serially diluted extract fractions in the microtiter plates. The plates were then checked for visible growth or turbidity to determine the MIC after 24 hours of incubation at 37°C. The lowest final well concentration of diluted extract fractions with no visible growth was regarded the MIC.

### Determination of minimum bactericidal concentration (MBC)

To confirm the MIC and determine the minimum bactericidal concentration (MBC), loopfuls of broth culture were taken from the wells that showed no detectable growth in the MIC test were streaked on fresh nutrient agar plates without the extract fractions. The plates were incubated at 37^0^C for 24 hours and examined for possible growth. Concentrations that did not show any growth after incubation were indicative of bactericidal effect and considered as the MBC [28].

## Data analysis

For dose-dependent analysis using the SYBR Green technique, percentage inhibition was determined by the formula, Percentage Inhibition = (1 – [RFU of treated-RFU of background] / [RFU of untreated – RFU of background] ×100%. For time-dependent analysis or kinetics, percentage parasitaemia was estimated by the formula, % Parasitaemia = iRBCs/tRBCs × 100, where iRBCs is the total number of infected red blood cells, and tRBCs is the total number of red blood cells in the same field. Growth inhibition was determined by comparing parasitaemia in treated samples to that of the positive control using the formula: [(z-a)/z] × 100%, where **z** is the percentage parasitaemia of the positive controls, and ***a*** is the percentage parasitaemia in the treatment group. The concentration of an extract needed to inhibit the growth the parasite by 50%. (IC₅₀) were also determined to assess the potency of each extract or fraction in inhibiting *Plasmodium falciparum*. For antibacterial activity, proportion of bacteria isolates inhibited the fractions were determined. Means and standard error of the means (SEM) were calculated. Where appropriate, statistical differences were assessed by analysis of variance (ANOVA) followed by multiple pairwise comparisons. A *p*-value less than 0.05 was considered statistically significant. All statistical analyses, including IC₅₀ determinations and dose-response curves, were performed using GraphPad Prism (GraphPad Software, San Diego, CA, USA).

## Results

### Antiplasmodial activity

#### Half-maximal inhibitory concentration of *Moringa oleifera* extracts against asexual *Plasmodium falciparum*

The antiplasmodial activity of *M. oleifera* seed extracts and fractions was evaluated using *in vitro* assays against the ring-stage of *P. falciparum* (3D7 strain). IC₅₀ values were determined to quantify the potency of each sample.

Among the crude extracts, the aqueous crude extract exhibited the highest inhibitory activity with an IC₅₀ of 107.0 µg/mL, followed by absolute ethanol at 160.0 ug/mL (Table 1). Among the extract fractions, the residual aqueous fraction had the best inhibitory activity at 135.3 µg/mL. Organic solvent fractions showed variable activity, with IC₅₀ values ranging from 398.2 to 696 µg/mL (Table 1). The IC_50_ of butanol could not be determined as its dose-dependent activity did not vary significantly.

**Table 1.** IC_50_ values for crude extracts and extract fractions of *Moringa oleifera* seed against asexual parasites.

| **Crude Extracts** | **IC<sub>50</sub> (95% IC)/ ug/ml** |
| --- | --- |
| Aqueous | 107 (85.1 to 141.5) |
| Absolute ethanol | 160.0 (73.6 to 184.6) |
| Petroleum ether | 494 (262.2 to 1108) |
| Ethyl acetate | 321.2 (NC) |
| **Extract fractions** |  |
| Hexane | 696 (357.2 to 2128) |
| Dichloromethane | 4101 (785.3 to 5216) |
| Ethyl acetate | 398.2 (286.4 to 658.7) |
| Butanol | ND |
| Residual | 135.3 (107.9 to 179.5) |
| **Control (Chloroquine)** | 0.0941 (0.0589 to 0.1618) |

NC: Not calculated. ND: Not determined

#### Dose-dependent variation in inhibition of parasite growth by extract fractions

The charts (Fig 2) showed a clear dose-dependent inhibition of *P. falciparum* asexual parasites across extract fractions. As the concentration decreased from 200 µg/mL to 3.12 µg/mL, the percentage inhibition reduced significantly for most of the extracts. The crude aqueous extract and residual fraction demonstrated the strongest activity, with inhibition exceeding 50% at 200 µg/mL. Whereas the percentage inhibition of crude aqueous extract and residual fraction did not differ significantly from that of chloroquine at 200 µg/mL (p > 0.05), the inhibitions of hexane, dichloromethane, ethyl acetate, and butanol were significantly lower than that of chloroquine at 200 µg/mL (p < 0.05). From 100 µg/mL to 3.12 µg/mL, there was a significant difference between chloroquine and all the extracts (p < 0.05). Generally, ethyl acetate had the lowest inhibition.

**Fig 2.**
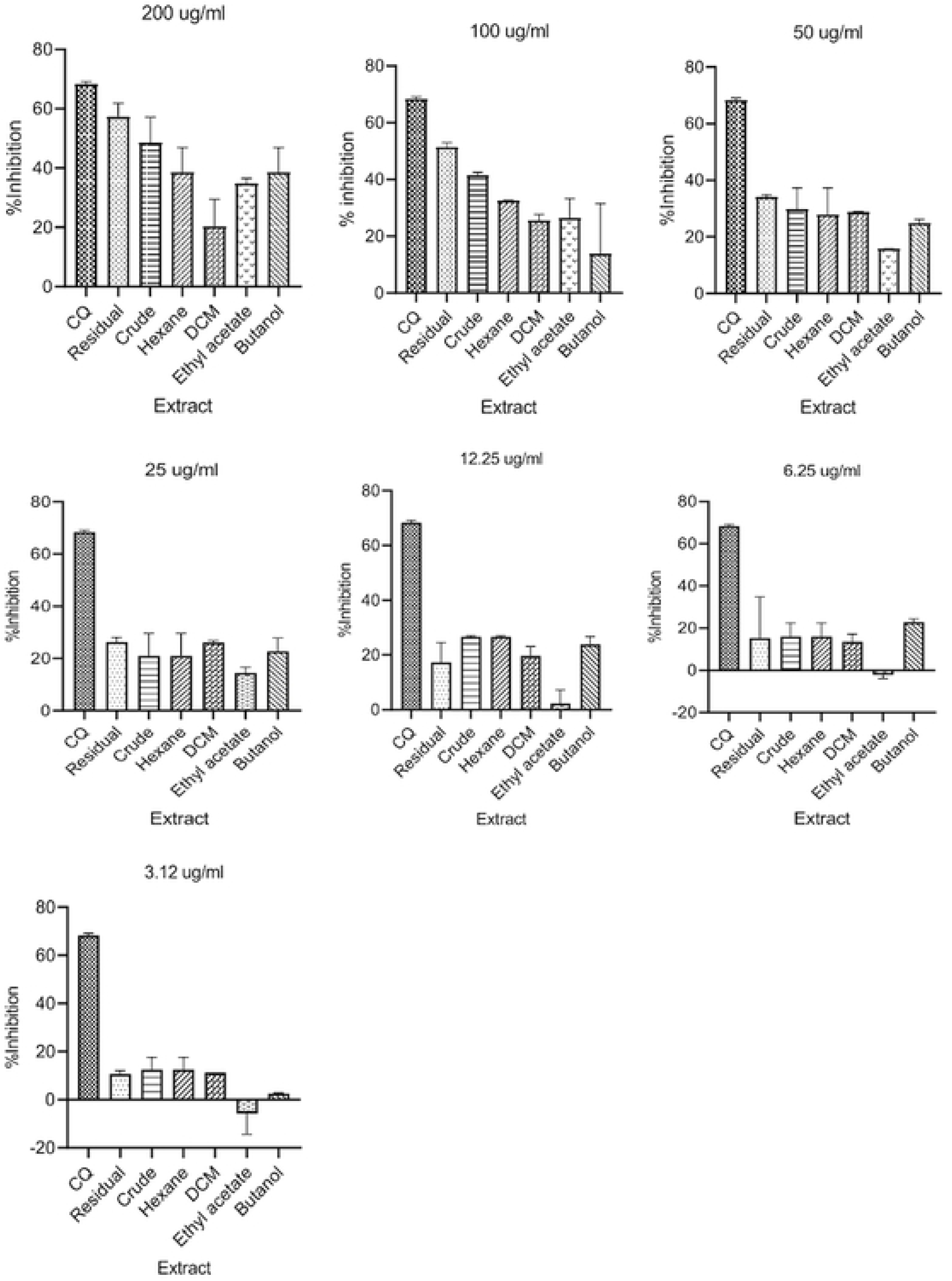
Dose-dependent inhibition *of parasite growth* following treatment with crude extract and extract fractions. The bars represent the median percentage inhibition and the range.

#### Time-dependent variation in percentage parasitaemia following treatment with crude extract and residual fraction at IC₅₀ and 2× IC₅₀ concentrations

In the analysis of time-dependent variation in percentage parasitaemia, as shown in the graph (Fig 3), a steady decline in percentage parasitaemia was observed over 48 hours when ring-stage parasites were treated with the crude extract, with 2× IC₅₀ concentration achieving faster and more pronounced parasite clearance compared to the IC₅₀. Similar to the crude extract, the residual fraction reduced % parasitaemia in a time-dependent manner, though with slightly delayed onset (Fig 4). Higher concentration (2 × IC₅₀) produced stronger activity.

**Fig 3.**
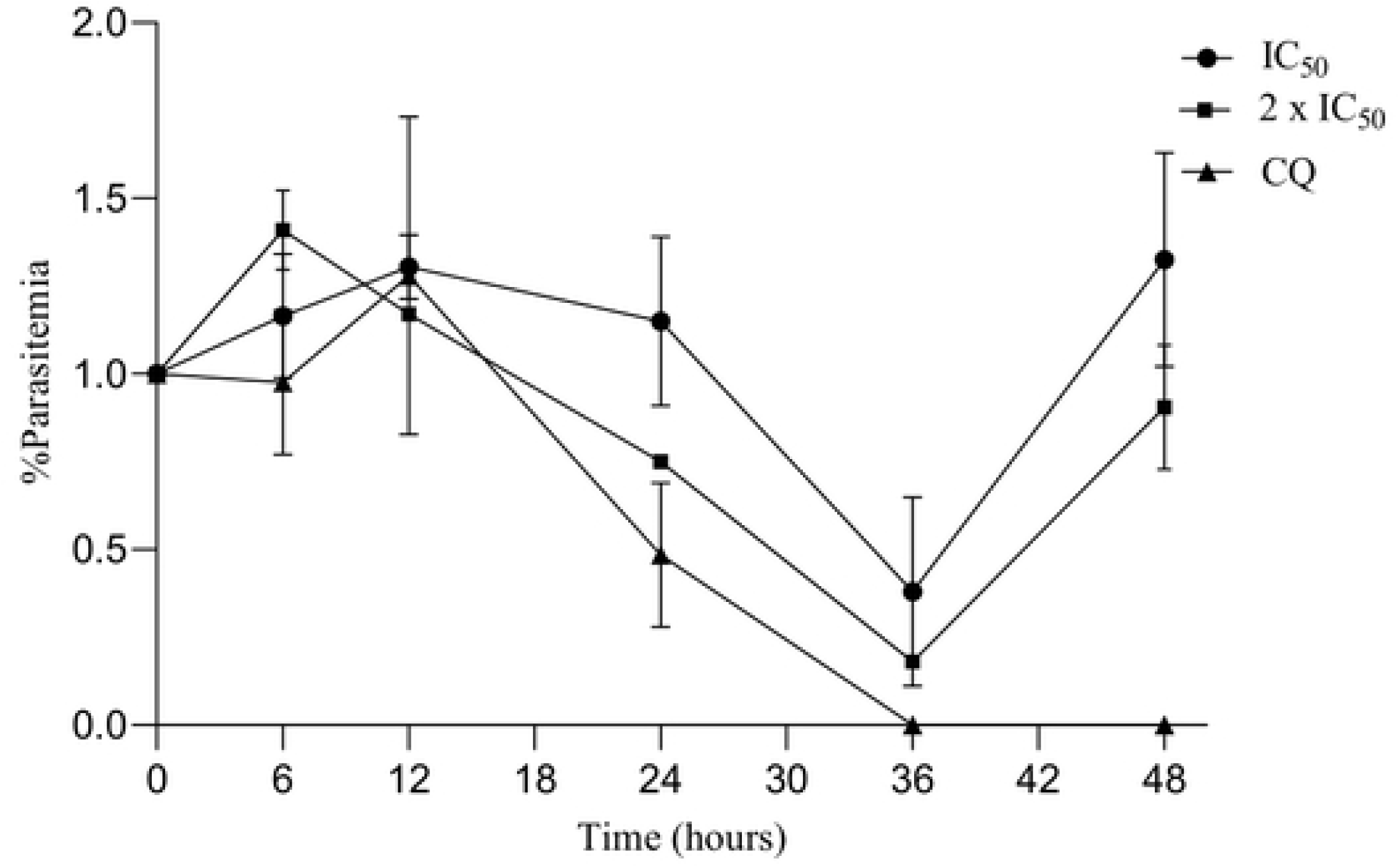
Time-dependent percentage parasitaemia with IC50 and 2 x IC50 concentration of crude extract. Assays were repeated three times; the points and error bars represent the median and range, respectively.

**Fig 4.**
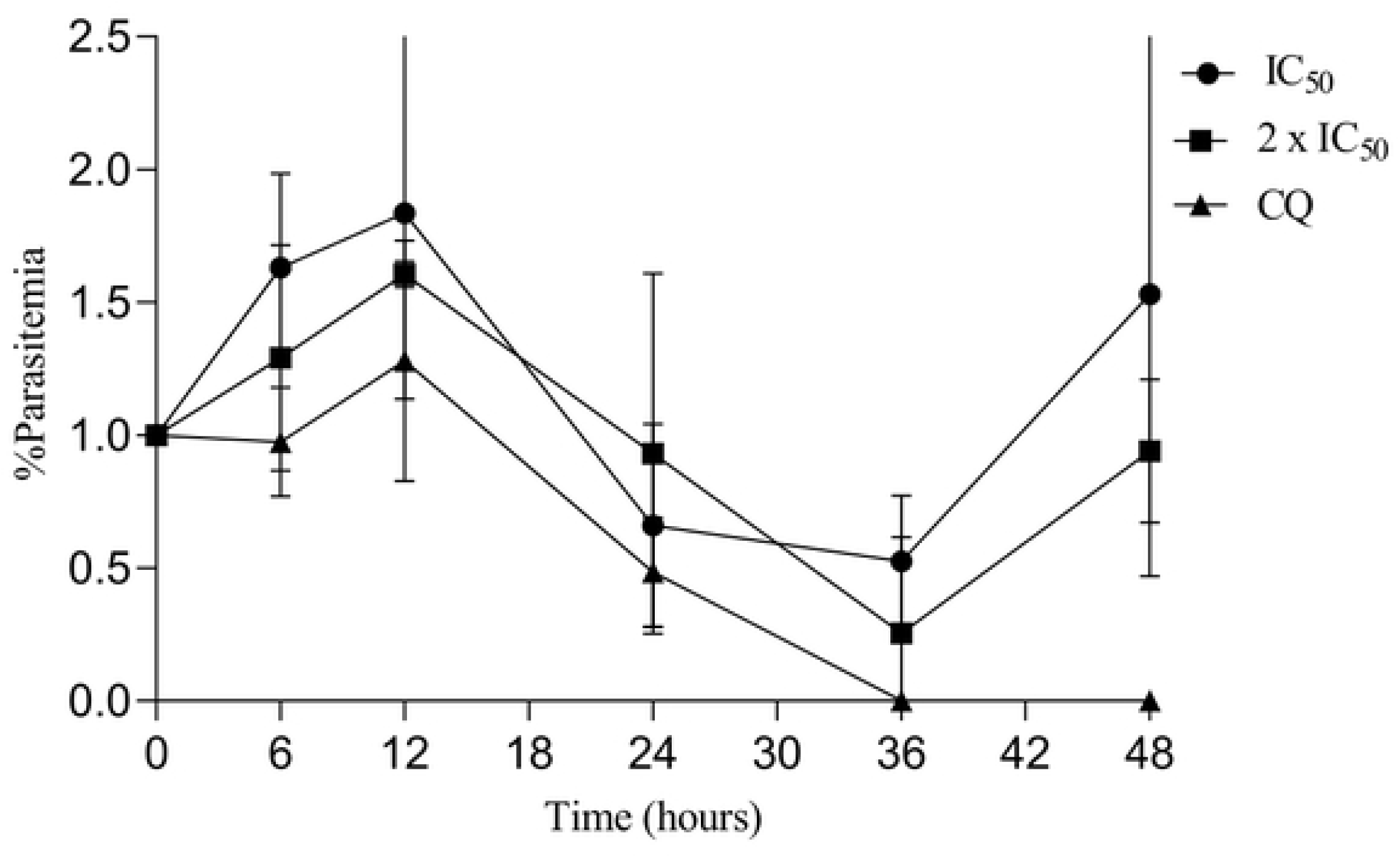
Time-dependent percentage parasitaemia with IC50 and 2 x IC50 concentration of residual fraction. Assays were repeated three times; the points and error bars represent the median and range, respectively.

Comparative analysis of crude extract and residual fraction at 2 × IC₅₀ concentration shows the superior efficacy of the residual fraction over the crude extract in reducing % parasitaemia over time (Fig 5). Both extracts showed substantial activity, though chloroquine remained the most potent. In all cases, % parasitaemia began to increase after 36 hours of incubation.

**Fig 5.**
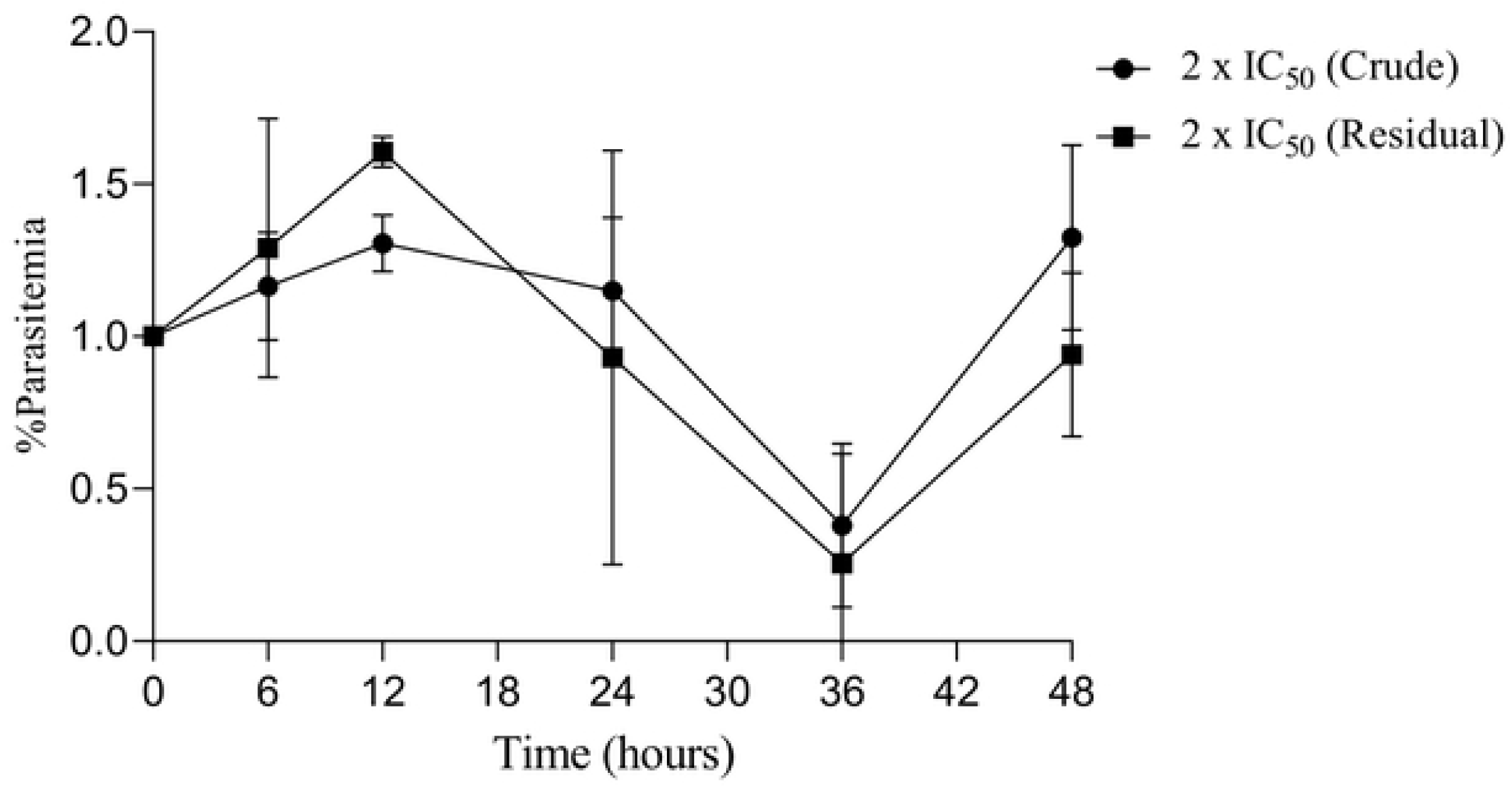
Time-dependent percentage parasitaemia with 2 x IC50 concentration of crude extract and residual fraction compared. Assays were repeated three times; the points and error bars represent the median and range, respectively.

#### Time-dependent variation in percentage inhibition of parasite growth by crude extract and residual fraction at IC₅₀ and 2 × IC₅₀ concentration

Crude extract and residual fraction demonstrated increasing % inhibition over time, reaching between 70–80% by 48 hours. Chloroquine maintained near-complete % inhibition throughout.

At higher concentrations (2 × IC₅₀), both extracts achieved >80% inhibition by 48 hours, and there were no significant differences between the percentage inhibition of the two extracts at both concentrations (Figs 6 and 7). Above 12 hours, the percentage inhibition of both extracts at either concentration (IC₅₀ or 2 × IC₅₀) did not differ from that of chloroquine (p>0.05).

**Fig 6.**
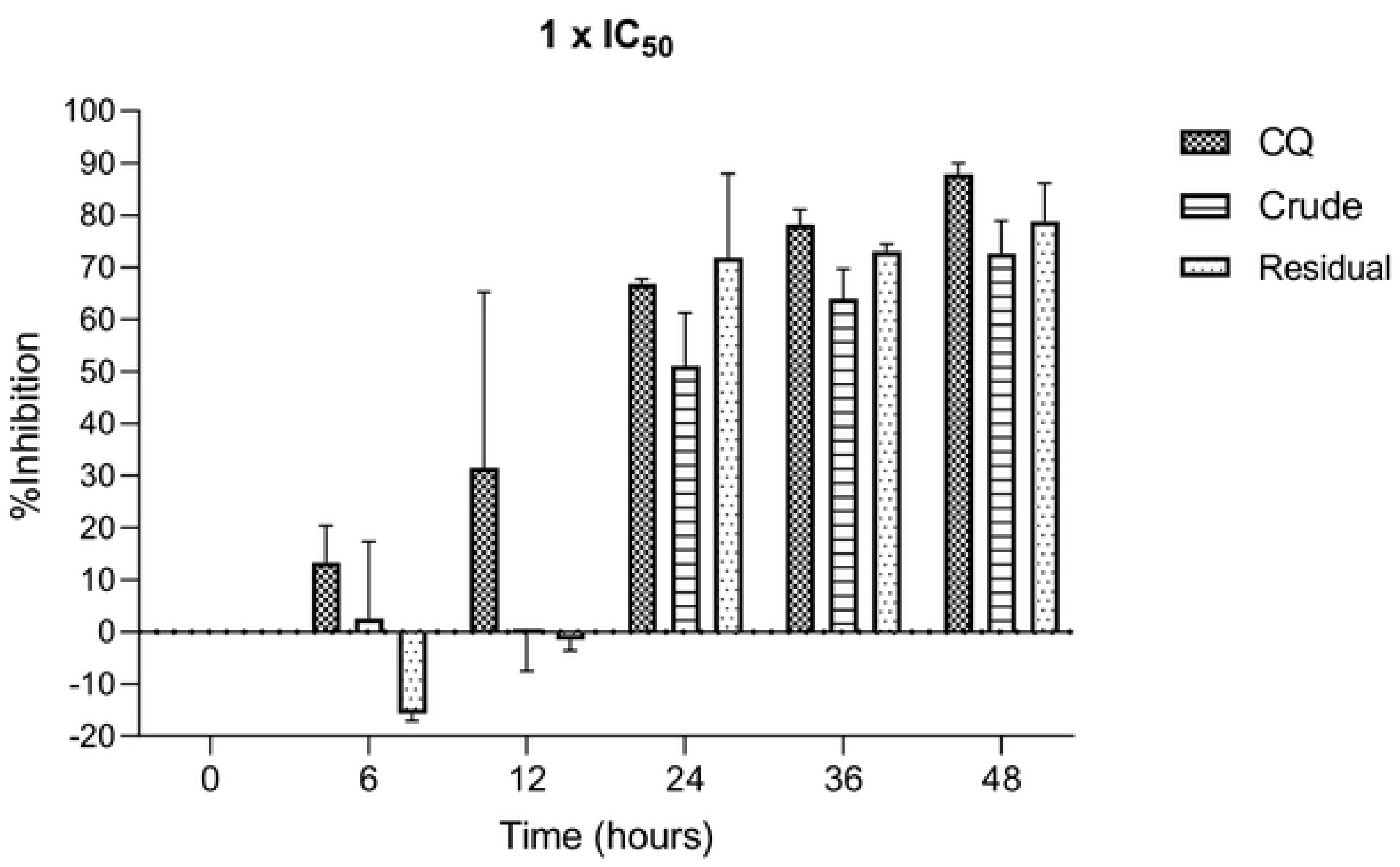
Time-dependent percentage inhibition by crude extract and residual fraction at IC₅₀ concentration. The bars represent the median percentage inhibition and the range.

**Fig 7.**
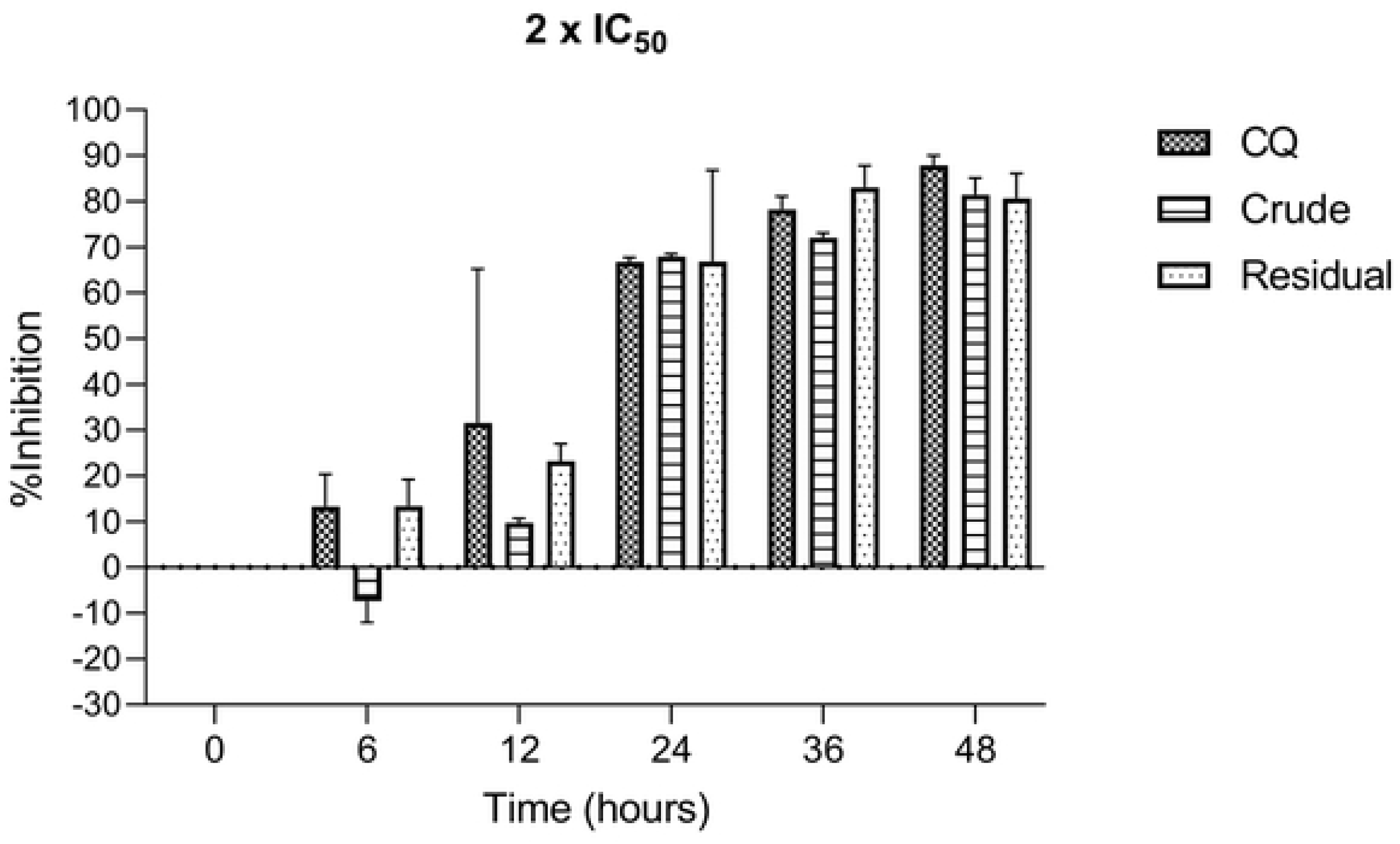
Time-dependent percentage inhibition by crude extract and residual fraction at 2 × IC₅₀ concentration. The bars represent the median percentage inhibition and the range.

### Antibacterial activity

#### Proportions of bacterial isolates inhibited by the extract fractions

Clinical and drug-resistant bacterial isolates were treated with 400µg/ml of fractions of *M. oleifera* seed extract and incubated for 24 hours. Various proportions of the isolates were inhibited by the fractions. The positive control (antibiotics) and ethyl acetate fraction inhibited all the isolates. All isolates of *S. aureus* (100%), 63% of *P. aeruginosa* and 88% of *E. coli* were inhibited by dichloromethane fraction. The ethyl acetate and dichloromethane fractions also inhibited drug-resistant isolates tested. The crude aqueous extract also inhibited most of the isolates, 80%, 50% and 88% for *S. aureus*, *P. aeruginosa* and *E. coli*, respectively. Hexane and butanol fractions inhibited fewer isolates than the ethyl acetate and dichloromethane fractions. The residual fraction showed no antibacterial activity (Table 2).

**Table 2:**
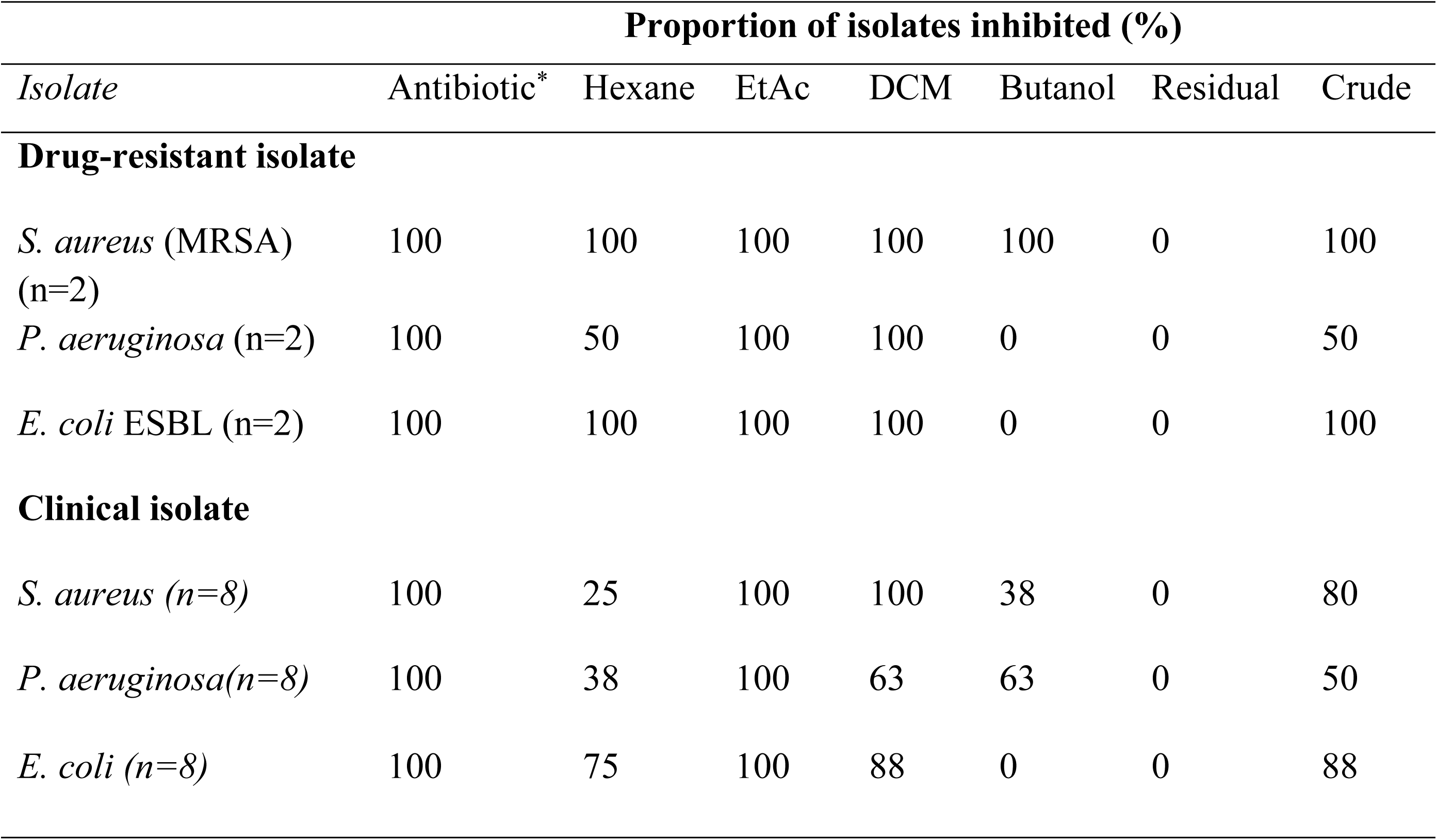
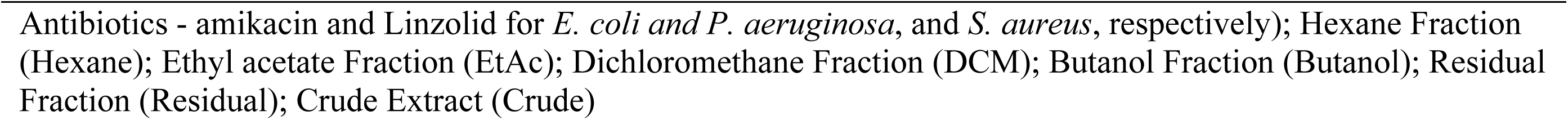
Proportion of bacteria isolates inhibited by 400 µg/ml of fractions of *M. oleifera* seed extract.

| Isolate | Proportion of isolates inhibited (%) |  |  |  |  |  |  |
| --- | --- | --- | --- | --- | --- | --- | --- |
|  | Antibiotic* | Hexane | EtAc | DCM | Butanol | Residual | Crude |
| **Drug-resistant isolate** |  |  |  |  |  |  |  |
| *S. aureus* (MRSA) (n=2) | 100 | 100 | 100 | 100 | 100 | 0 | 100 |
| *P. aeruginosa* (n=2) | 100 | 50 | 100 | 100 | 0 | 0 | 50 |
| *E. coli* ESBL (n=2) | 100 | 100 | 100 | 100 | 0 | 0 | 100 |
| **Clinical isolate** |  |  |  |  |  |  |  |
| *S. aureus* (n=8) | 100 | 25 | 100 | 100 | 38 | 0 | 80 |
| *P. aeruginosa* (n=8) | 100 | 38 | 100 | 63 | 63 | 0 | 50 |
| *E. coli* (n=8) | 100 | 75 | 100 | 88 | 0 | 0 | 88 |

Antibiotics - amikacin and Linzolid for *E. coli* and *P. aeruginosa*, and *S. aureus*, respectively); Hexane Fraction (Hexane); Ethyl acetate Fraction (EtAc); Dichloromethane Fraction (DCM); Butanol Fraction (Butanol); Residual Fraction (Residual); Crude Extract (Crude)

#### Inhibitory activity of fractions compared

Ethyl acetate and dichloromethane fractions had the best activity against all the isolates. For the clinical isolates, analysis of variance (ANOVA) showed significant variations in the zones of inhibition of the fractions for all the isolates; *S. aureus* (p< 0.0001), *P. aeruginosa* (p=0.0004) and *E. coli* (p< 0.0001).

Pairwise comparisons revealed that the zone of inhibition of ethyl acetate and dichloromethane fractions for *S. aureus* were significantly higher than those of hexane fraction, butanol fraction and crude extract (p<0.05), while no significant difference existed between the zones of inhibition of ethyl acetate and dichloromethane fractions. The crude aqueous extract also had a significantly higher zone of inhibition compared the hexane fraction against *S. aureus*.

For *P. aeruginosa*, ethyl acetate again had significantly higher zones of inhibition compared hexane fraction, butanol fraction and crude extract (p<0.05), while no significant variation existed between the zones of inhibition of ethyl acetate and dichloromethane fractions. The zones of inhibition of *E. coli* by ethyl acetate fraction was also higher than those of hexane and butanol fractions (p<0.05). Butanol also had significantly lower zone of inhibition than those of dichloromethane fraction, hexane fraction and crude extract (p<0.05). Similar trend was observed for the resistant and the control strains (Fig 8).

**Fig 8.**
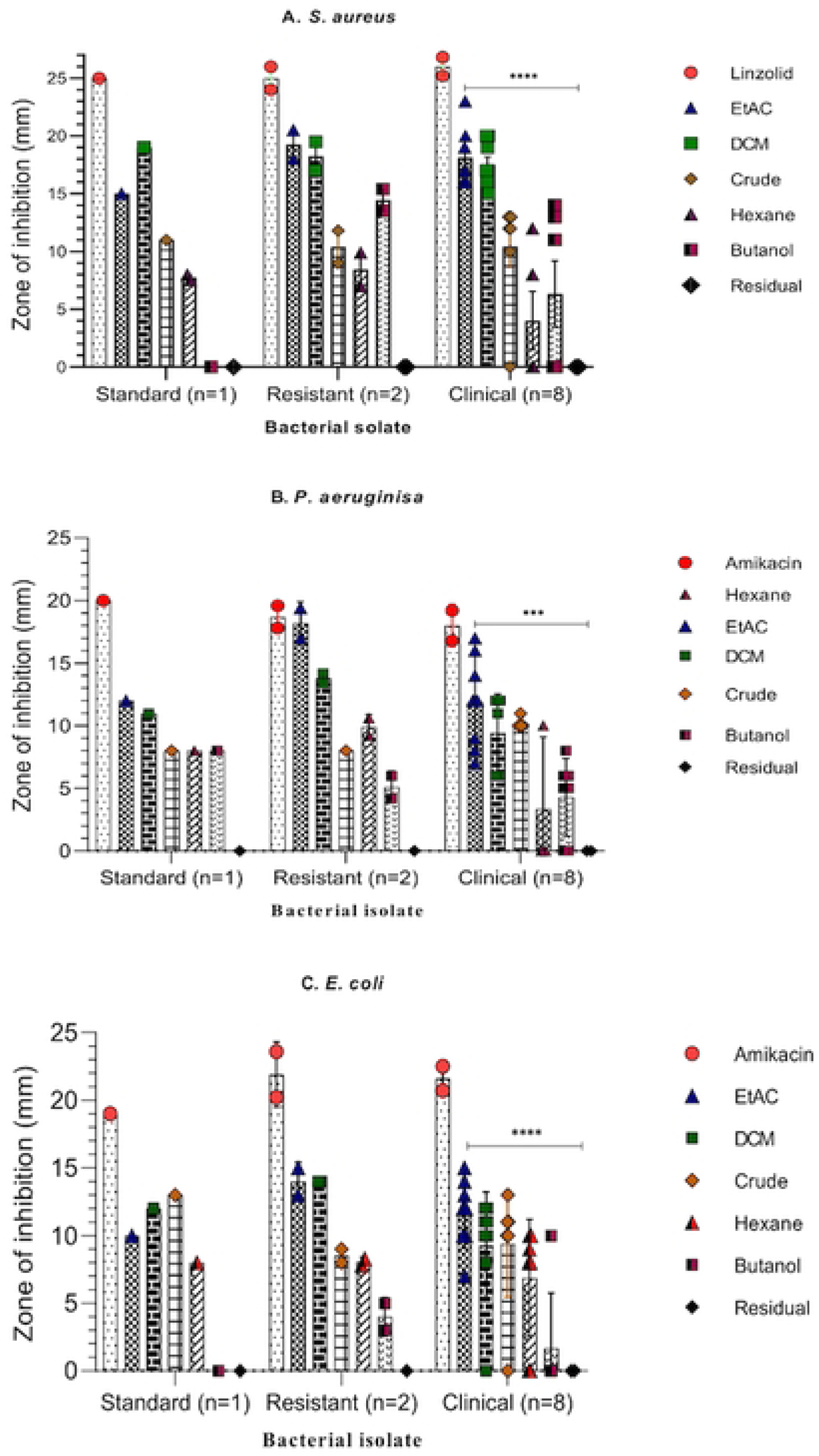
Zone of inhibition for bacteria isolates tested with 400µg/ml of fractions of *M. oleifera* seed extract. *S. aureus* isolates (A); *P. aeruginosa* isolates (B); C. *E. coli* isolates; Antibiotics - amikacin was used as positive control for *E. coli* and *P. aeruginosa*, and Linzolid for *S. aureus*); hexane fraction (Hexane); ethyl acetate fraction (EtAc); dichloromethane fraction (DCM); butanol fraction (butanol); residual fraction (Residual); crude extract (Crude), S. aureus resistant isolates were methicillin-resistant *S. aureus* (MRSA). *E. coli* resistant isolates were extended spectrum β-Lactamase (ESBL)-producing *E. coli*. Where applicable, the values are the mean ± SEM of parallel measurements. ***: p< 0.001, and ****: p< 0.0001.

#### Minimum inhibitory (MIC) and minimum bactericidal (MBC) concentrations

The values for the MIC and MBC of the fractions and the crude extract against the isolates of *S. aureus, P. aeruginosa* and *E. coli* are presented in Table 3. The MICs varied depending on the fraction and isolate of bacteria examined. The MICs ranged from 6.3 to 12.5µg/ml for ethyl acetate, 25 to 50 µg/ml for dichloromethane and 50 µg/ml for the crude extract. The MBC ranged from 25 to 100 µg/ml for ethyl acetate fraction, 50 to 100 µg/ml for dichloromethane and 100 µg/ml for the crude extract. It is interesting to note that both the MICs and MBCs were consistently lower for ethyl acetate fraction than for both dichloromethane fraction and crude extract. However, the MBC/MIC ratio for all the extracts was ≤4, indicating universal bactericidal activity (Table 3).

**Table 3:** Minimum inhibitory concentration (MIC) and minimum bactericidal concentration (MBC) for ethyl acetate and dichloromethane fractions on isolates of *S. aureus, P. aeruginosa* and *E. coli*.

| MIC (MBC) of Fractions (µg/ml) |  |  |  |  |  |  |
| --- | --- | --- | --- | --- | --- | --- |
| Isolate | EtAc |  | DCM |  | Crude |  |
|  | MIC<br>(MBC) | MBC/MIC<br>ratio | MIC<br>(MBC) | MBC/MIC<br>ratio | MIC<br>(MBC) | MBC/MIC<br>ratio |
| **Drug-resistant Isolate** |  |  |  |  |  |  |
| *S. aureus* (1) | 25 (100) | 4 | 25 (50) | 2 | 50 (100) | 2 |
| *S. aureus* (2) | 25 (50) | 2 | 25 (50) | 2 | 50 (100) | 2 |
| *P. aeruginosa* (1) | 25 (50) | 2 | 50 (100) |  | 50 (100) | 2 |
| *P. aeruginosa* (2) | 25 (50) | 2 | 50 (100) | 2 | 50 (100) | 2 |
| *E. coli* (1) | 12.5 (25) | 2 | 25 (50) | 2 | 50 (100) | 2 |
| *E. coli* (2) | 12.5 (25) | 2 | 25 (50) | 2 | 50 (100) | 2 |
| **Clinical isolates** |  |  |  |  |  |  |
| *S. aureus* (1) | 6.25 (25) | 4 | 25 (50) | 2 | 50 (100) | 2 |
| *S. aureus* (2) | 6.25 (25) | 4 | 50 (100) | 2 | 50 (100) | 2 |
| *P. aeruginosa* (1) | 25 (50) | 2 | 50 (100) | 2 | 50 (100) | 2 |
| *P. aeruginosa* (2) | 25 (50) | 2 | 50 (100) | 2 | 50 (100) | 2 |
| *E. coli* (1) | 50 (100) | 2 | 25 (50) | 2 | 50 (100) | 2 |
| *E. coli* (2) | 50 (100) | 2 | 25 (50) | 2 | 50 (100) | 2 |
| Ethyl acetate Fraction (EtAc); Dichloromethane Fraction (DCM); Crude Extract (Crude). |  |  |  |  |  |  |

## Discussion

*Moringa oleifera* leaves and seeds have been used for treating malaria in many countries [12,18], and their antiplasmodial potential has been demonstrated in some studies [12, 21,22]. Likewise, studies have also demonstrated that aqueous and organic solvent extracts of *M. oleifera* seeds and leaves exhibit inhibitory effects on both enteric and non-enteric microbes [11,23,24]. However, the antiplasmodial properties of *M. oleifera*, particularly the seeds, and its effectiveness in treating malaria and bacterial infections have not been fully unravelled.

The present study demonstrates that *M. oleifera* seed extracts possess measurable antiplasmodial activity against the chloroquine-sensitive *P. falciparum* 3D7 strain, with aqueous crude extracts and residual aqueous fractions showing the most promising inhibitory effects. The aqueous crude extract showed the highest potency among extracts, although, its activity remains moderate compared with chloroquine (IC₅₀ = 0.0941 µg/mL), indicating a large therapeutic gap relative to standard antimalarial drugs per the classification proposed by Rasoanaivo et al. [29], that IC₅₀ values between 50–200 µg/mL be considered to have moderate activity. The dose-dependent inhibition observed in this study further supports the therapeutic potential of *M. oleifera*. At concentrations of 200 µg/mL, both crude aqueous extract and residual fraction achieved inhibition levels comparable to chloroquine, although activity declined significantly at lower concentrations. This pattern is consistent with many studies that demonstrated that *M. oleifera* extracts exhibited dose-dependent suppression of *Plasmodium* growth (17).

The relatively high IC₅₀ values in this study indicate that while *M. oleifera* seed extracts show promise, they are less potent than other plant extracts [30,31] and established antimalarials and may require optimization through purification of active compounds. Even if *Moringa* seeds extract may unlikely yield highly potent standalone antimalarials, it may serve as a source of supportive agents or lead compounds for combination therapies. The moderate activity is still significant in the context of rising resistance to artemisinin-based therapies, as adjunctive plant-derived compounds could help delay resistance development [4].

Kinetic assays revealing time-dependent inhibition of development or growth indicated that both crude aqueous extract and residual fraction reduced parasitaemia in a time-dependent manner, achieving >80% inhibition at 48 hours, particularly at 2× IC₅₀ concentrations. This gradual clearance mirrors the delayed but sustained activity reported for other plant-derived antimalarials, such as *Azadirachta indica* (neem) extracts [32].

A decline in parasitaemia up to approximately 36 hours followed by rebound is consistent with partial suppression during trophozoite/schizont maturation and subsequent re-invasion when drug pressure is insufficient or wanes [33]. Plant extracts often act more slowly or with narrower stage windows, explaining the delayed onset and the stronger effect at 2× IC₅₀ [34; 35]. The faster decline at 2× IC₅₀ fits classic pharmacodynamic expectations where higher exposure increases killing rate and reduces viable schizont output [33]. The rise after 36 hours suggests either surviving rings progressing to schizonts and re-invasion, reduced effective concentration, or instability of bioactive substances over time, which are common in crude botanical assays [33].

The fractionation process brought new clarity by separating activities according to solvent polarity. The aqueous crude extract exhibited an IC₅₀ of 107 µg/mL, while the residual aqueous fraction recorded 135.3 µg/mL, both significantly lower than the IC₅₀ values of organic solvent fractions such as hexane, dichloromethane, and ethyl acetate. Though the high IC₅₀ values indicate weak antiplasmodial activity, the findings suggest that highly polar compounds retained in aqueous extracts may be responsible for the observed activity. This aligns with earlier reports that polar phytochemicals, including alkaloids, flavonoids, and glycosides, often contribute to antimalarial efficacy [36, 14], among others, through interference with haeme polymerization and parasite metabolic pathways as is the case with flavonoid [37].

Antibacterial activity of the fractions was also examined. The ethyl acetate and dichloromethane fractions exhibited potent antibacterial activity, inhibiting all isolates of *S. aureus* (100%) and *E. coli* (88-100%), and a significant proportion of *P. aeruginosa* isolates (63-100%). These findings are consistent with previous studies that reported the antibacterial activity of *M. oleifera* extracts against various bacterial pathogens [14, 23, 24, 38]. The crude aqueous extract also demonstrated notable antibacterial activity, inhibiting 80% of S. aureus, 50% of *P. aeruginosa*, and approximately 88% of *E. coli* isolates. Analysis of variance revealed statistically significant differences in zones of inhibition (*p* < 0.0001 for *S. aureus* and *E. coli*), confirming the superior efficacy of ethyl acetate and dichloromethane fractions. Pairwise comparisons, however, indicated no significant difference between ethyl acetate and dichloromethane fractions, but both were superior to hexane, butanol, and crude extracts. This result is consistent with our previous observation that crude ethyl acetate extract had higher zones of inhibition compared to other organic extracts [25, 39]. The MIC and MBC values further supported the potent antibacterial activity of the ethyl acetate fraction, with lower MICs (6.3-12.5 µg/ml) and MBCs (25-100 µg/ml) compared to the dichloromethane fraction and crude extract. Importantly, the MBC/MIC ratio of ≤4 indicated that the extracts exhibited bactericidal activity against the tested isolates [40; 41] rather than bacteriostatic activity Interestingly, the strong antibacterial activity against MRSA and ESBL-producing *E. coli* is particularly significant in the context of global antimicrobial resistance (AMR) [42]. Plant-derived phenolics and isothiocyanates have demonstrated membrane-disruptive and enzyme-inhibitory mechanisms against resistant bacteria [14] and this could be the case with *M. oleifera* anti-bacterial activity.

In general, the polarity of extraction solvents played a decisive role in shaping the biological activity of the fractions. Polar aqueous extracts (crude and residual fractions) exhibited the strongest antiplasmodial activity, with IC₅₀ values of 107–135 µg/mL, while moderately polar organic fractions or semi-polar fractions (ethyl acetate and dichloromethane) showed superior antibacterial activity. Non-polar fractions (hexane) were generally weak in both assays, indicating that depending on the solvent for the extraction, the use of *M. oleifera* in treating fevers mediated by malaria parasite or bacteria may not be effective, especially in traditional settings. However, the use of the whole plant part, for instance, the seed, may be efficacious. Though ethyl acetate was the second to last solvent in the fractionation process, it captured most active substances, indicating that most of the active substances have lower polarity but not as low as that of hexane. Fractionation thus revealed the possible classes of compounds are responsible for the two activities, and importantly, it clarified that the residual aqueous fraction retained antiplasmodial potency while losing antibacterial activity. This separation of bioactivity is a novel finding, as many earlier studies relied on crude extracts without fractionation, which obscured the specific contributions of solvent polarity [26]. Meanwhile, future work should focus on bioassay-guided fractionation, structural characterization of active molecules, and *in vivo* validation to determine pharmacological relevance and safety.

## Acknowledgements

The technical support of Isaac Junior Okyere, Grace Ocloo Semevor of the Department of Medical Microbiology, University of Ghana Medical School, and Jessica Asomaniwaa Armah of the Department of Chemistry, University of Ghana, is duly acknowledged.

